# GWAS significance thresholds in large cohorts

**DOI:** 10.1101/2024.12.09.627629

**Authors:** Evans K Cheruiyot, Tingyan Yang, Allan F McRae

## Abstract

While the p-value threshold of 5.0 × 10^−8^ remains the standard for genome-wide association studies (GWAS) in humans and other species, it still needs to be updated to reflect the current era of large-scale GWAS, where tens of thousands of sample sizes are used to discover genetic associations at loci with smaller minor allele frequencies. In this study, we used a dataset of 348,501 individuals of European ancestry from the UK Biobank to determine the GWAS thresholds required for multiple testing corrections when considering rare and common variants in additive and dominant GWAS models. Additionally, we employed conditional and joint (COJO) analysis to quantify the proportion of false significant hits in the GWAS results for 72 traits in the UK Biobank when applying the traditional GWAS cut-off versus our newly proposed p-value thresholds. Overall, the results indicate that the conventional GWAS significance threshold of 5.0 × 10^−8^ yields a false positive rate of between 20% and 30% in GWAS studies that utilize large sample sizes and less common variants. Instead, a more stringent GWAS p-value threshold of 5.0 × 10^−9^ is needed when rare variants (with minor allele frequency > 0.1%) are included in the association test for both additive and dominance models within the European ancestry population.

## Introduction

The widely accepted p-value threshold for genome-wide association studies (GWAS) of common variants in humans is 5.0 × 10^−8^. This threshold was developed based on smaller sample sizes and marker densities available during the early days of GWAS (International Hapmap Consortium: 2005; Dudbridge AND Gusnanto 2008) and has been highly successful in discovering many reproducible genetic variants associated with common traits and diseases. However, two remarkable advances in the GWAS era necessitate revisiting and updating this p-value threshold. First, the sample sizes used in GWAS have increased steadily over the years, reaching over 5 million individuals in recent studies (Yengo *et al*. 2022). Second, with the advances in large-scale genotyping and next-generation sequencing technologies, GWA studies have focused not only on common variants (MAF > 5%) but also on low-frequency (1% < MAF < 5%) and rare (MAF < 1%) variants, which have been shown to contribute substantially to complex trait heritability (Lee *et al*. 2014).

Several studies have investigated GWAS significance threshold in the context of deep phenotyping and wide allele frequency spectrum in GWAS (Xu *et al*. 2014; Fadista *et al*. 2016; Kanai *et al*. 2016; Asif *et al*. 2021; Chen *et al*. 2021). However, these studies have used relatively small sample sizes in their simulations to derive GWAS thresholds and are thus outdated when performing GWAS in large biobank cohorts. In addition, large sample sizes are providing power to investigate non-additive modes of association. In particular, the traditional GWAS traditional cut-off (5.0 × 10^−8^) has also been used for dominance models (Palmer *et al*. 2023; Zhu *et al*. 2023). However, different GWAS thresholds for additive and dominance models would be expected since the extent of linkage disequilibrium (LD) tagging across variants differs between the two models (Zhu *et al*. 2023).

Various statistical approaches have been used to determine GWAS significance thresholds: the Bonferroni correction (which assumes independent tests and thus overly conservative considering that the genetic variants are in LD) (Risch AND Merikangas 1996), permutation and bootstrapping (resampling) (International Hapmap Consortium: 2005; Kanai *et al*. 2016; Lin 2019), and Bayesian methods (Wellcome Trust Case Control 2007). Other methods involving calculating the ‘effective number of tests’ based on LD pruning or eigenvalue decomposition of phenotypes and genotypes (Cheverud 2001; Duggal *et al*. 2008; Sobota *et al*. 2015) have also been proposed. However, while the permutation-based method is computationally demanding, it remains a “gold standard” in GWAS multiple testing correction because it preserves correlation structure for the data and is robust to the assumption of marker independence (Dudbridge AND Gusnanto 2008; Hoggart *et al*. 2008; Asif *et al*. 2021).

## Results and discussion

We used a permutation procedure to estimate the GWAS significance thresholds using the UK Biobank (version 3) dataset (Bycroft *et al*. 2018). We restricted our analyses to a cohort of 348,501 unrelated individuals with European ancestry. Our results revealed that a more stringent GWAS cut-off than the commonly used threshold (5.0 × 10^−8^) is required to control false-positive rate if variants with MAF of less than 5% are included in the association test (Table 1). The traditional GWAS p-value cut-off of 5.0 × 10^−8^ is still valid when considering common (>5% MAF) variants in the additive GWA model using large sample sizes. However, a stringent GWA p-value threshold of 8.9 × 10^−9^ is needed when rare variants (MAF > 0.1%) are included in an additive GWA model (Figure 1; Table 1). A more stringent GWA correction is required to guard against false positives when the phenotype is not normally distributed. This was demonstrated using our BMI based trait, where the p-value cut-off was determined to be 5.9 × 10^−9^ for MAF > 0.1% (Figure 1). Using the traditional threshold of 5.0 × 10^−8^ leads to substantial Type-I error, particularly with low MAF thresholds (Table 2). When rare variants (MAF > 0.1%) are included in the association tests, we observed a false positive rate of approximately 22% for a perfectly normal trait in the additive model, increasing to 30% with some phenotype skewness.

**Table 1.** Estimated GWAS significance thresholds for different minor allele frequency cut-off obtained from additive and dominance models.

| Model | MAF cut-off* | P<sub>sig</sub> threshold | M<sub>eff</sub> |
| --- | --- | --- | --- |
| Additive GWA (Normal) | > 0.1% | $8.9 \times 10^{-9}$ | 5,636,978 |
| | > 0.5% | $1.7 \times 10^{-8}$ | 2,873,563 |
| | > 1% | $2.8 \times 10^{-8}$ | 1,785,714 |
| | > 5% | $4.9 \times 10^{-8}$ | 1,014,198 |
| Additive GWA (Raw BMI) | > 0.1% | $5.9 \times 10^{-9}$ | 8,417,508 |
| | > 0.5% | $1.5 \times 10^{-8}$ | 3,378,378 |
| | > 1% | $1.7 \times 10^{-8}$ | 2,873,563 |
| | > 5% | $3.3 \times 10^{-8}$ | 1,497,005 |
| Dominance (Normal) | > 0.1% | $7.4 \times 10^{-9}$ | 6,720,430 |
| | > 0.5% | $1.5 \times 10^{-8}$ | 3,246,753 |
| | > 1% | $1.9 \times 10^{-8}$ | 2,564,102 |
| | > 5% | $3.5 \times 10^{-8}$ | 1,404,494 |
| Dominance (Raw BMI) | > 0.1% | $5.7 \times 10^{-8}$ | 8,756,567 |
| | > 0.5% | $1.3 \times 10^{-8}$ | 3,937,007 |
| | > 1% | $1.4 \times 10^{-8}$ | 3,597,122 |
| | > 5% | $2.7 \times 10^{-8}$ | 1,818,181 |

**Table 2.** Proportion of associations that passed traditional threshold (5.0 × 10^−8^) but failed to reach the updated threshold for the additive GWAs.

| Model | Trait | MAF cut-off |  |  |  |
| --- | --- | --- | --- | --- | --- |
|  |  | > 0.1% | > 0.5% | > 1% | > 5% |
| Additive GWA | Simulated normal | 22.2 | 13.1 | 9.5 | 5.1 |
|  | Raw BMI | 29.5 | 16.6 | 13.3 | 7.3 |

**Figure 1.**
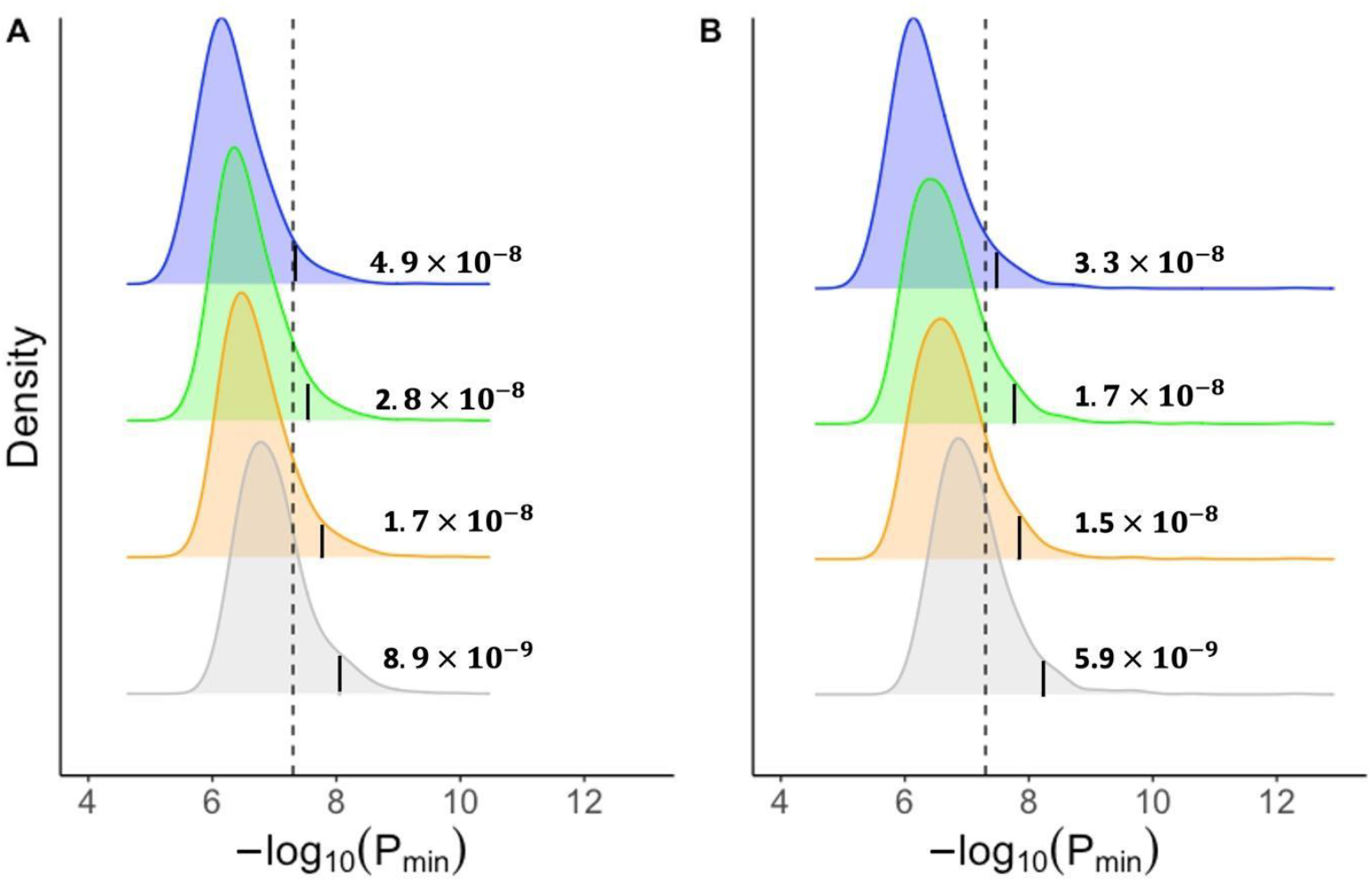
The distribution of the minimum p-value (−*log*_10_*P*_*min*_) from the additive GWAS models. We extracted −*log*_10_*P*_*min*_ from a GWA results for each of the (**A**) 1,000 simulated normally distributed phenotypes and (**B**) permuted raw body mass index (BMI) phenotypes from UK Biobank. The genotypes used in GWA were from the European cohort in the UK Biobank (N = 348,501). The dotted line corresponds to the traditional genome-wide significance cut-off of (5.0 × 10^−8^), while the vertical solid lines represent the new empirical GWAS cut-off after pruning genotypes based on the > 5% (blue), >1% (green), > 0.5% (orange), and > 0.1% (grey) minor allele thresholds. The new cut-off GWAS thresholds represents 95% percentile of *P*_min_ distribution (α = 0.05).

These findings are consistent with previous studies showing that a more stringent p-value cut-off is required when the MAF pruning is relaxed to accommodate rare variants in GWAS (Fadista *et al*. 2016). Our results are comparable with simulations using data (∼ 140k) from European ancestry that obtained a GWA significance p-value of 6.6 × 10^−9^ for MAF ≥ 0.1% (Kemp *et al*. 2017). This indicates that for MAF > 0.1%, the effect of increasing sample size is attenuated by the time sample sizes reach hundreds of thousands (Asif *et al*. 2021). As such, our point estimates are likely applicable even for large GWAs (Yengo *et al*. 2022). Taken together, we recommend a more stringent GWAS p-value cut-off of 5.0 × 10^−9^ if rare variants (MAF > 0.1%) are considered in the association test for an additive model – equivalent to 10 million independent tests at 5% alpha (Table 1).

We also investigated the p-value threshold when applying a dominance model in GWAS (Figure 2). It is common for gene mapping studies to assume a typical GWA p-value (5.0 × 10^−8^) for both additive model additive and dominance GWAS, particularly for common variants^10^. However, a more stringent threshold is needed to minimise false positive results under the dominance model – even for common variants – because dominance LD tagging captures less variation than additive LD tagging in the genome^11^. Our results show that a lower p-value threshold of 3.5 × 10^−8^ is needed when mapping common variants (MAF > 5%) in a dominance GWAS (Figure 2). Even a smaller p-value cut-off is ideal if the phenotype is skewed, as shown from our simulations based on the UK Biobank BMI trait where we found a p-value threshold of 2.7 × 10^−8^ (Figure 2 and Table 1). When rare variants (MAF > 0.1%) are included the GWAS, we found a p-value cut-off of 7.4 × 10^−9^ (for normally distributed trait and a 5.7 × 10^−9^ for a slightly skewed phenotypes (Figure 2 and Table 1).

**Figure 2.**
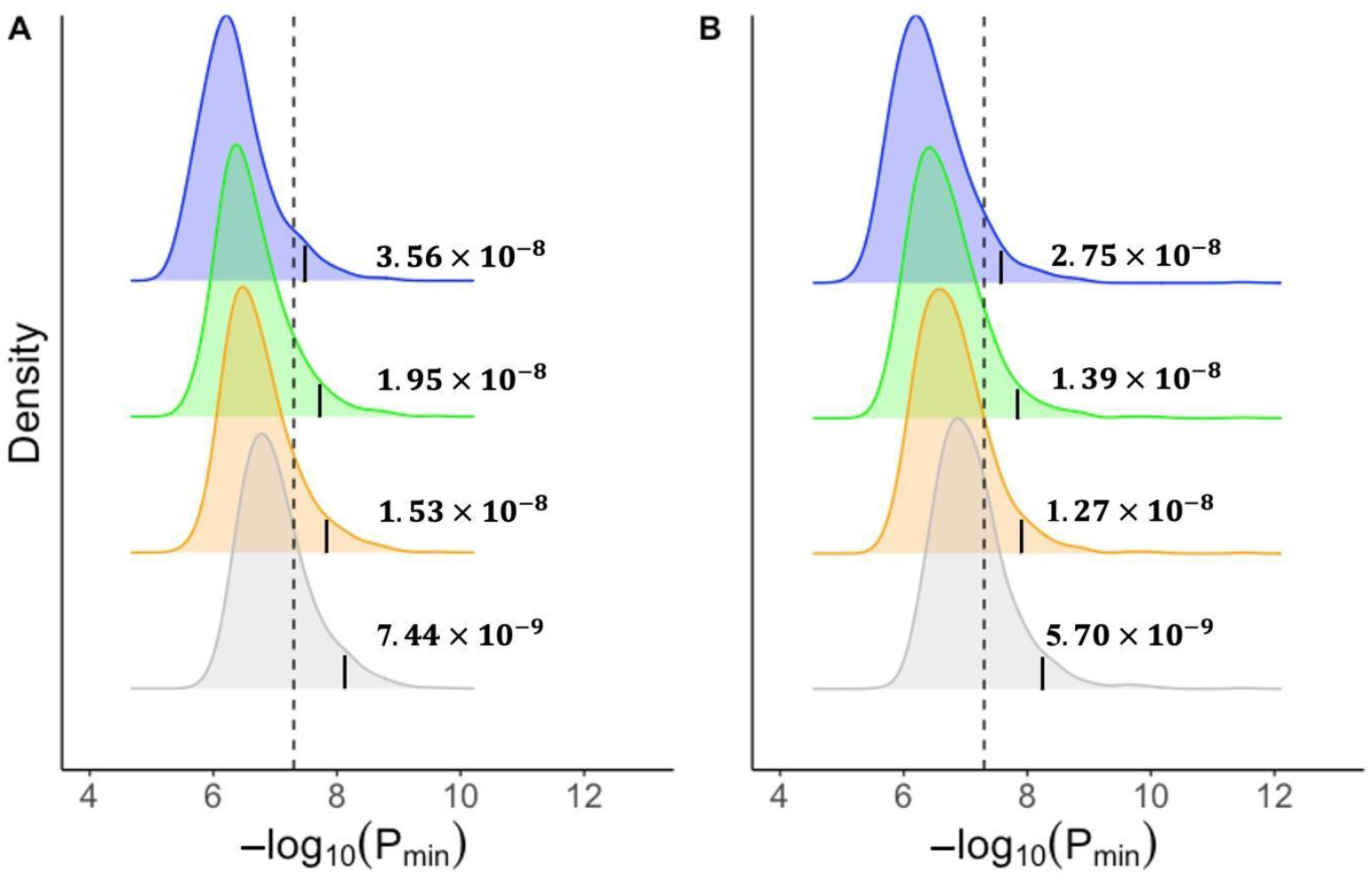
The distribution of minimum p-value (−*log*_10_*P*_*min*_) from the dominance GWAS models. The −*log*_10_*P*_*min*_ was extracted from a GWA results based on the (**A**) 1,000 simulated normally distributed phenotypes and (**B**) permuted raw body mass index (BMI) phenotypes from UK Biobank. The genotypes used were from the European cohort in the UK Biobank (N = 348,501). The dotted line corresponds to the traditional genome-wide significance cut-off of (5.0 × 10^−8^), while the vertical solid lines represent the new empirical GWAS cut-off after pruning genotypes based on the > 5% (blue), > 1% (green), > 0.5% (orange), and > 0.1% (grey) minor allele thresholds. The new cut-off GWAS thresholds represents 95% percentile of *P*_min_ distribution (α = 0.05).

We used conditional and joint (COJO) analysis (Yang *et al*. 2012) implemented in the GCTA software (Yang *et al*. 2011) to identify independent associations and quantify the proportion of tests in the GWA results from UK Biobank that passed traditional significance threshold (5.0 × 10^−8^) but failed to reach the new empirical GWAS threshold from our simulations – referred hereafter as the “false significance rate (FSR)”. Here, we quantified the FSR from the most stringent empirical GWAS thresholds obtained from filtering genotypes based on the MAF cut-off of > 0.1% for only the additive GWAS simulations. COJO performs conditional analysis of GWA summary statistics and does not require individual-level genotypes. To run COJO, we downloaded GWA summary statistics for 72 quantitative traits in the UK Biobank (GWAS round 2). The summary statistics for the raw and inverse-rank normalized (irnt) phenotypes were available for these traits. We used a random subset of 10,000 unrelated Europeans from the UK Biobank as linkage disequilibrium (LD) reference in the COJO analysis. We ran COJO per chromosome for the GWA summary data for both sexes using the default setting. We calculated the FSR for each trait as the percentage difference in the number of independent GWAS counts between the traditional GWA significance threshold (5.0 × 10^−8^) and the new empirical GWA threshold.

By using the most stringent empirical GWA correction from the additive model based on the MAF > 0.1% cut-off (8.9 × 10^−9^), we found the proportion of false significant results for the inverse normalized traits ranged from 5.56% (Glycated haemoglobin (HbA1c)) to 51.43% (microalbumin in urine), with an average of 18.26% across traits (Figure 3 and Supplementary Table S1). Similarly, applying the updated p-value correction from the simulations for the raw phenotypes (5.9 × 10^−9^) in COJO, yielded a slightly higher average proportion of false significant results of 20% across non-normalized traits. Besides microalbumin in urine, the other normalized traits with the highest false significance rate included creatinine and sodium in urine and fluid intelligence score (Figure 3). We found similar levels of FSR when using the software plink “clumping” approach (with the default parameters) instead of COJO, with the average false significant results across traits of 20.95% and 23.29% for normalized (irnt) and non-normalized (raw) traits, respectively (Supplementary Table S2). Given the move to large meta-analysis leaving few suitable cohorts with power for replication, these results highlight the need for properly controlled type-I error to avoid follow-up studies on potential false-positive associations.

**Figure 3.**
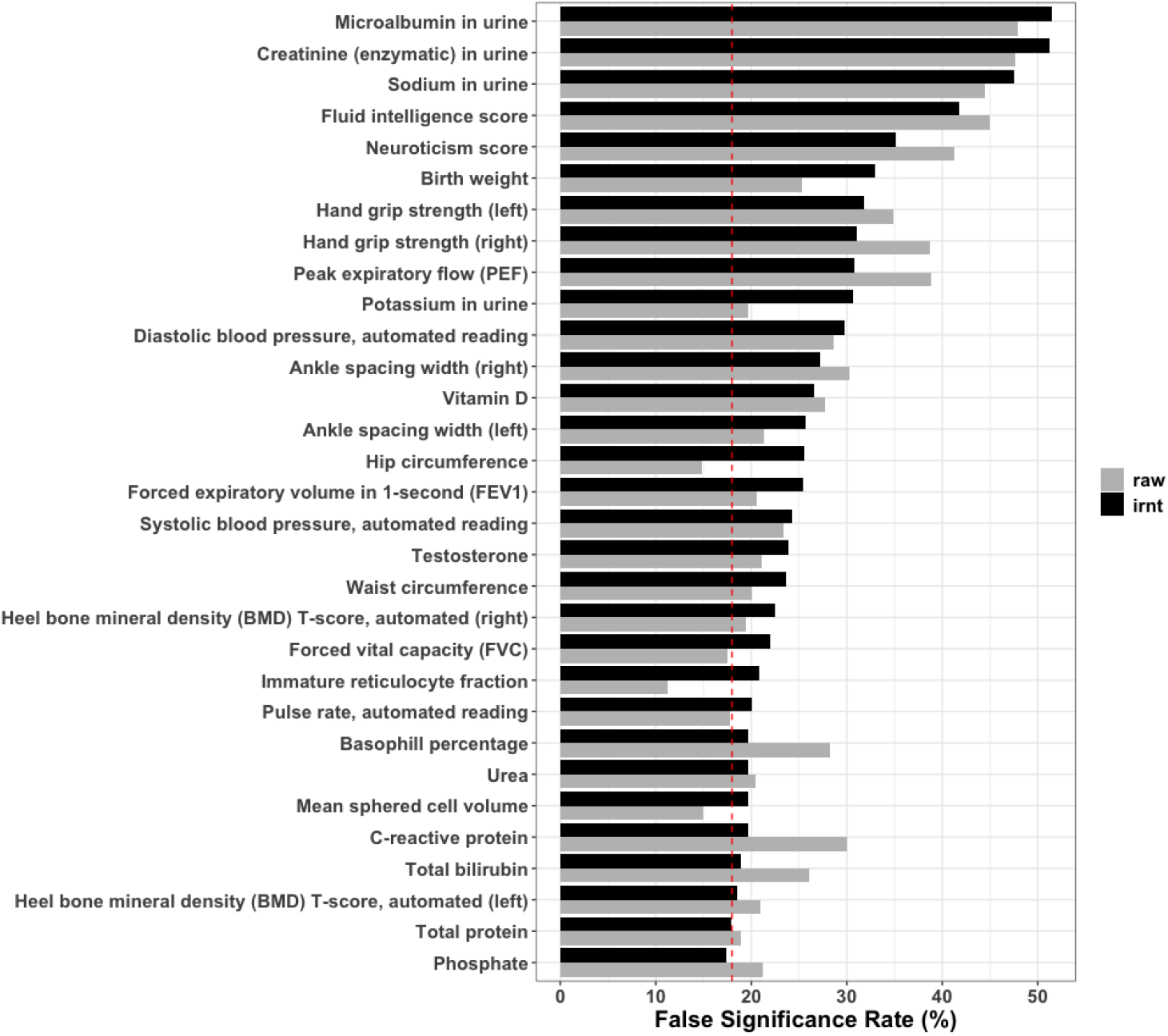
Top 30 traits with the highest proportion of associations (‘false significance rate’ – FSR) that passed traditional threshold (5.0 × 10^−8^) but failed to reach the updated threshold for the additive GWAS. The FSR for the inverse rank normalized transformed (irnt) traits and their corresponding raw untransformed phenotypes are represented by black and grey colors, respectively. The FSR was calculated as the percentage of minimum p-values (−*log*_10_*P*_*min*_) passing the typical GWAS cut-off (5.0 × 10^−8^) from the GWAS of the 1,000 simulated traits. The red vertical dashed line represents average FSR for the irnt traits.

Notably, our empirical GWA threshold applies to only the European ancestry population, which we used in the simulations because a large sample size was available for GWAS. Future studies on other populations, e.g., Africans, are required given that LD structure is population specific and, therefore, different GWAS correction thresholds are needed (Kanai *et al*. 2016; Pulit *et al*. 2017).

In summary, using simulations with large sample sizes, we have established that the traditional GWAS significance p-value of 5.0 × 10^−8^ is insufficient for multiple testing correction in current GWAS that use large sample sizes and less common variants. Instead, a stringent GWAS correction p-value of 5.0 × 10^−9^ is needed when rare variants (MAF > 0.1%) are considered in the association test for both additive and dominance models in the European ancestry population. Adopting the new p-value threshold for rare variants is critical for guarding against type 1 errors in future GWAS for European ancestry.

## Declaration of interests

The authors declare no competing interests.

## Acknowledgments

Allan McRae is the recipient of an Australian Research Council Australian Fellowship (project number FT200100837) funded by the Australian Government. The views expressed herein are those of the authors and are not necessarily those of the Australian Government or Australian Research Council. This research has been conducted using the UK Biobank Resource under project 12505.

## Author contributions

A.F.M conceived and designed the study. E.K.C, T.Y and A.F.M conducted data analysis.

E.K.C wrote the manuscript. All authors reviewed, revised and approved the final manuscript for publication.

## Web Resources

Uk Biobank GWAS data, https://www.nealelab.is/uk-biobank

## Data and code availability

This work used genotype and phenotype data from UK Biobank Resource under project 12505. UKB data can be accessed upon request once a research project has been submitted and approved by the UKB committee.

## Materials and methods

We simulated two sets of 1,000 phenotypes were reflecting: 1) the ideal statistical scenario where the phenotype is normally distributed (or has undergone rank-based inverse normal transformation) and 2) a trait with skewness. To simulate skewness, we used body mass index (BMI) measurements from the UK Biobank participants (regardless of inclusion in the genetic data). BMI data was standardized within sex to have a mean of zero and unit variance and values with scaled BMI > 6 (∼5 inter-quartile range) were removed. Then, we randomly sampled the scaled BMI without replacement to generate 1000 independent traits.

To estimate GWA significance thresholds across different minor allele frequency (MAF) scenarios, the genotype data was pruned based on four MAF cut-offs [> 0.1%, > 0.5%, > 1% and > 5%]. A total of 13,035,294, 9,603,317, 8,545,459, and 6,098,223 autosomal variants remained following MAF filtering at these thresholds, respectively.

For each of the simulated phenotypes [N = 1,000 x 2 models], we ran additive and dominance GWAS analysis. We used the *fastGWA* method implemented in GCTA software [option *--fastGWA-lr*] (Yang *et al*. 2011) to run additive linear models, while the dominance GWAS was performed using PLINK version 2 software [*--glm dominant*] (Purcell *et al*. 2007). No covariates were included in all the GWAS models.

To identify genome-wide significance thresholds for each of the MAF cut-offs, we examined the distribution of the minimum p-values −*log*_10_*P*_*min*_ for each simulated trait, separately for additive and dominance models. The 95% percentile of the −*log*_10_*P*_*min*_ distribution was defined as the genome-wide significance threshold for each MAF filtering scenario, reflecting a 5% genome-wide significance p-value. The false positive rate when using the traditional GWAS significance threshold for each scenario was calculated as the proportion of traits for which −*log*_10_*P*_*min*_ surpassed the genome-wide significance threshold of 5.0 × 10^−8^.

